# OptiJ: Open-source optical projection tomography of large organ samples

**DOI:** 10.1101/656488

**Authors:** Pedro P. Vallejo Ramirez, Joseph Zammit, Oliver Vanderpoorten, Fergus Riche, François-Xavier Blé, Xiao-Hong Zhou, Bogdan Spiridon, Christopher Valentine, Simeon P. Spasov, Pelumi W. Oluwasanya, Gemma Goodfellow, Marcus J. Fantham, Omid Siddiqui, Farah Alimagham, Miranda Robbins, Andrew Stretton, Dimitrios Simatos, Oliver Hadeler, Eric J. Rees, Florian Ströhl, Romain F. Laine, Clemens F. Kaminski

**Affiliations:** Laser Analytics Group, Department of Chemical Engineering and Biotechnology, University of Cambridge, Cambridge UK; Sensor CDT 2015-2016 student cohort, University of Cambridge, Cambridge, UK; Clinical Discovery Unit, Early Clinical Development, IMED Biotech Unit, AstraZeneca, Cambridge, UK; Bioscience, Respiratory, Inflammation and Autoimmunity, IMED Biotech Unit, AstraZeneca, Gothenburg, Sweden; Medical Research Council Laboratory for Molecular Cell Biology (LMCB), University College London, Gower Street, London, WC1E 6BT; Department of Physics and Technology, UiT The Arctic University of Norway, NO-9037 Tromsø, Norway

**Keywords:** 3D imaging, OPT, projection tomography, lungs, open-source, low-cost, large organs, ImageJ, Fiji

## Abstract

The three-dimensional imaging of mesoscopic samples with Optical Projection Tomography (OPT) has become a powerful tool for biomedical phenotyping studies. OPT uses visible light to visualize the 3D morphology of large transparent samples. To enable a wider application of OPT, we present OptiJ, a low-cost, fully open-source OPT system capable of imaging large transparent specimens up to 13 mm tall and 8 mm deep with 50 μm resolution. OptiJ is based on off-the-shelf, easy-to-assemble optical components and an ImageJ plugin library for OPT data reconstruction. The software includes novel correction routines for uneven illumination and sample jitter in addition to CPU/GPU accelerated reconstruction for large datasets. We demonstrate the use of OptiJ to image and reconstruct cleared lung lobes from adult mice. We provide a detailed set of instructions to set up and use the OptiJ framework. Our hardware and software design are modular and easy to implement, allowing for further open microscopy developments for imaging large organ samples.

## Introduction

The three-dimensional imaging of anatomical and functional features in mesoscopic biological samples (millimeter-scale dimensions) e.g. in model organisms, organs, or even plants, provides valuable data for biomedical research. Standard 3D imaging techniques such as micro-MRI (1–4) and micro-CT (5–9) are used in biomedical imaging to visualize morphology in large tissues and organs at micrometer-level resolution. However, these techniques are expensive and cannot take advantage of molecule-specific labeling strategies that are available to fluorescence microscopy. Confocal (10) or light sheet fluorescence microscopy (11–13) can be used to generate volumetric data with optical sectioning at sub-cellular resolution, although the usable specimen sizes are typically confined to sub-millimeter scales and commercial microscopy systems can be expensive. Optical Projection Tomography (OPT)(14) is a 3D imaging technique for transparent mesoscopic samples which allows visualizing micrometer-scale features. OPT is based on computerized tomography techniques (15) in which 2D images, called projections, are acquired with different sample orientations and then used to obtain a 3D image of the sample using a reconstruction algorithm, such as filtered-back projection (FBP). Sample clearing is often necessary to allow light propagation and imaging through the thickness of the sample. OPT can operate using either absorption/scattering of the sample (transmission OPT, tOPT) or fluorescence (emission OPT, eOPT) to generate image contrast. The use of OPT has been reported widely, and applications include the visualization of the 3D anatomy of mouse embryos (16–27), zebrafish (21, 24, 28–34), drosophila (35–38), plants (39, 40), *C.elegans* (41), animal organs (22, 27, 42–44) and other mesoscopic samples (45–47). Although major improvements in resolution (48, 49), acquisition time (31), field of view (FOV) (21, 40) and compatibility with other imaging techniques (22, 28, 50) have been made, most OPT applications require advanced technical expertise, expensive equipment, and bespoke software for reconstruction.

To enable a more general uptake of this technique, we present OptiJ (Fig. 1a-b), a low-cost, integrated, open-source implementation of OPT specifically designed to enable the 3D imaging of large organ samples in both fluorescence and transmission modes. Our framework includes a complete set of open-source ImageJ/Fiji (51) plugins to reconstruct OPT data from specimens up to 13 mm tall and 8 mm deep (13×8×8 mm^3^). A number of algorithms were developed to improve image quality. We include a thorough description of how to build and operate the hardware and how to use the software. Other open-source OPT implementations have been demonstrated for smaller volumes than what is necessary for large murine organs (24, 52), or for large volumes using commercial reconstruction software (21). Here, we demonstrate the capabilities of OptiJ by imaging full-sized adult mouse lungs that have been cleared and immunostained. Their study is relevant in the context of chronic obstructive pulmonary diseases (COPDs), which are characterized by heterogeneously distributed emphysema (alveolar cell death) and bronchoconstriction (narrowing of airways). OptiJ allowed us to explore the morphology of the airway tree and visualize in 3D the tertiary airways, bronchioles, and alveolar sacs in complete murine lungs. We share our results using FPBioimage (53), an open-source online visualization tool, so that readers can view and explore the reconstructed OPT data interactively in any standard web browser.

**Fig. 1.**
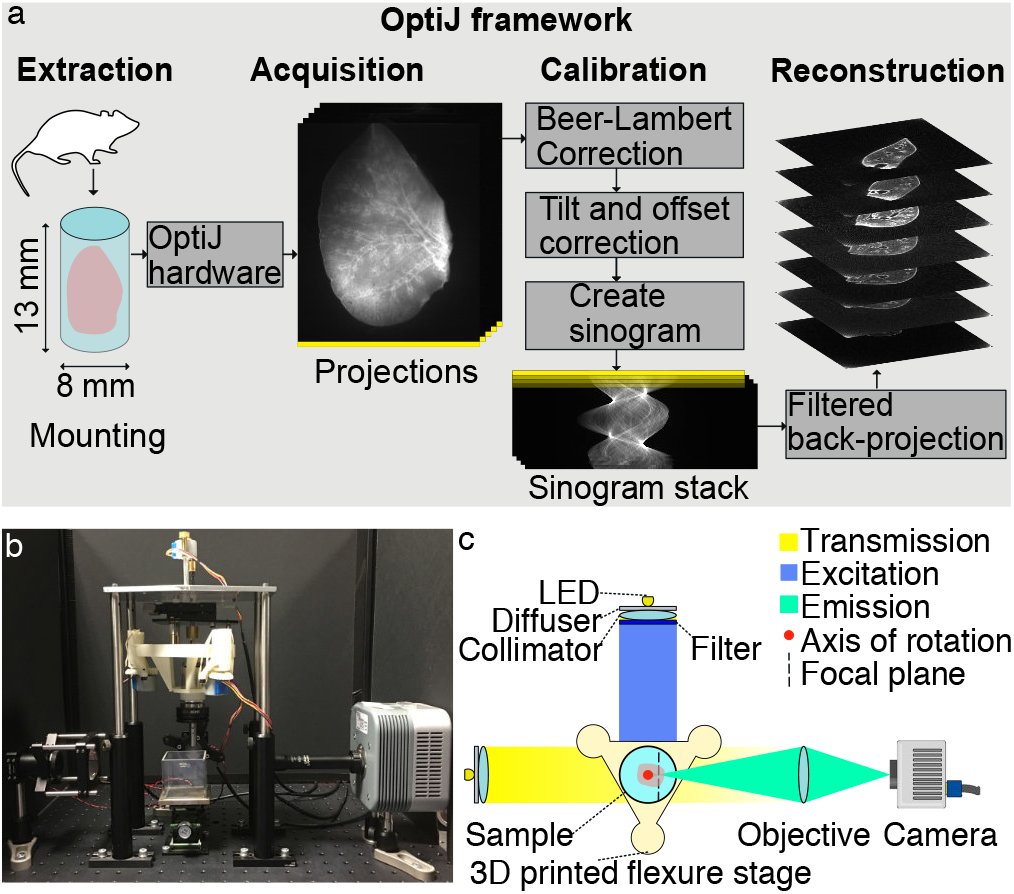
Schematic representation of the OptiJ Framework. **a)** OptiJ workflow including sample mounting, acquisition of projections, correction, and reconstruction steps. **b)** Picture of the OptiJ set-up. **c)** Top-view illustration of the OptiJ hardware.

## Results

### OptiJ hardware

The OPT principle relies on the rotation of a sample to acquire 2D projections at different angles. Assuming the thickness of the sample is less than the depth of field of the system, projections acquired over half a revolution are theoretically sufficient to recover an accurate 3D reconstruction of the sample structure. However, a full revolution typically leads to higher image quality (14, 31). When implementing our OptiJ system, we focused on the following considerations: (1) ensuring that the axis of rotation is parallel to the imaging plane of the camera, (2) aligning the sample to the field of view of the camera, and (3) robustly and repeatably performing the rotation of the sample and acquisition of the projections. The OptiJ hardware enables the mounting, alignment, and rotation of thick biological samples for the acquisition of 2D projections in both eOPT and tOPT modalities. Figure 1b-c shows the implemented set up, which includes a monolithic 3D-printed rotation and translation stage, a telecentric relay lens, a camera, two broadband LEDs, fluorescence excitation and emission filters, and collimating and diffusing optics. The main criteria guiding our component choice were ease of access, widespread availability, and low cost. The 3D-printed stage is adapted from the published Flexscope design (54) to accomplish the movement necessary for both linear alignment and rotation of the sample with low-cost stepper motors. The stage achieves sub-micron steps, with a maximal hysteresis of 58 μm over a 3 mm travel range (see Supplementary Information for details on the stage characterization). A low numerical aperture (NA) 0.5x telecentric lens was chosen to match the typical volume of adult mouse lungs. The low NA allows a depth of field of ~4 mm, which upon sample rotation allows for a maximum field of view of 13×8×8 mm^3^. The focal plane of the objective is placed midway between the axis of rotation and the front face of the sample such that only one half of the sample is in focus at any given projection angle (as shown with the dashed line in Fig.1 c). The telecentricity of the lens allows us to use the highly efficient FBP reconstruction approach. LEDs emitting over a broad spectral range were chosen for their brightness and long life, and a custom circuit board was designed to minimize output flicker. The LED output was homogenized and collimated with off-the-shelf optics to ensure uniform illumination across the field of view. The stage, the camera, and the LEDs were controlled with a RaspberryPi™ that interfaces with a central computer. A detailed description of the OptiJ hardware assembly, parts list, and system characterization can be found in the Supplementary Information.

### OptiJ analysis

The reconstruction of a high-quality 3D volume from the OPT projections requires data pre-processing to avoid artifacts during reconstruction via FBP. OptiJ includes a set of freely available ImageJ/Fiji plugins to preprocess OPT data, as well as an efficient GPU-enabled FBP algorithm for reconstruction. The plugins and the suggested workflow for their use is shown in Fig. 2a. The Beer-Lambert correction plugin divides each tOPT projection by an average brightfield image following the Beer-Lambert Law(55) to obtain linear attenuation coefficients corrected for non-uniform pixel intensities, as demonstrated in the lower panel of Fig. 2b.i. A common artifact in OPT arises from the axis of rotation of the sample not being parallel to the plane of the FOV during acquisitions, which leads to the appearance of a shadow artifact around sharp features as demonstrated in Fig. 2b.ii. The Estimate Tilt and Offset plugin tracks a fiducial marker (such as a 100 μm glass bead) in the projections to determine if the axis of rotation is parallel to the center of the FOV, and produces correction values for the projection stack if this condition is not satisfied. These values can be used at the reconstruction step to minimize any shadow artifacts, as demonstrated in the corrected image in Fig. 2.b.ii. The Create Sinogram plugin displays a Radon Transform of the projections and uses the correction values for tilt and offset produced by the previous plugin to account for residual deviations, relaxing the need for thoroughly precise alignment of the system prior to acquisitions. The output of this plugin is a sinogram, an intermediate step in the FBP reconstruction named after its sinusoidal shape. Small sample wobble caused by mechanical jitter from low-cost stepper motors can be detected as jagged edges in an otherwise smooth sinogram, demonstrated in Fig. 2b.iii. The Dynamic Offset Correction plugin calculates a sinusoidal fit of the motion of a fiducial marker and uses the difference between the ideal fit coordinates and the actual motion of the bead to produce a jitter-free sinogram as shown in the corrected image in Fig. 2b.iii. This step concludes the pre-processing required to minimize artifacts prior to reconstruction. The 2D reconstruction plugin implements an FBP algorithm to reconstruct a 3D cross-sectional stack of the original object using the corrected sinogram. To speed up reconstruction times via FBP, the plugin allows for GPU-enabled acceleration using OpenCL (56), which is open-source and platform independent. This plugin also allows the user to choose from a variety of filters (Ramp, Hamming, Shepp-Logan, or no filter) for back-projection (15). A detailed description of the OptiJ plugin library, its functions and methods, usage and sample data for testing can be found in our online repository at https://lag-opt.github.io.

**Fig. 2.**
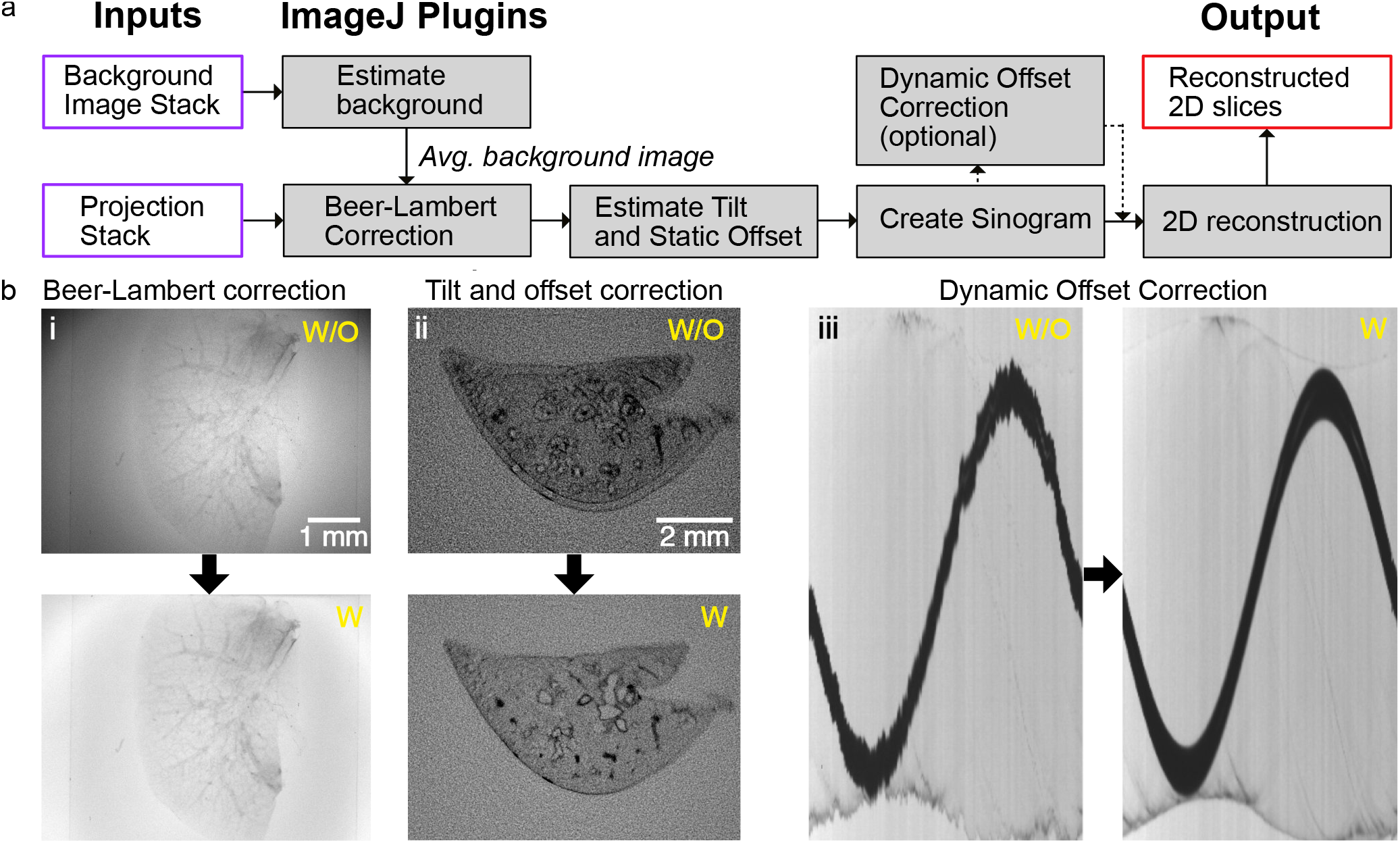
OptiJ plugin library workflow for the correction of common OPT artifacts. **a)** Typical workflow for the use of the OptiJ plugins. **b)** Correction of common OPT artifacts using OptiJ plugins. The top row represents images without correction applied (w/o). The bottom row shows images after correction (w). (i) Uneven illumination in raw tOPT projections resulting from the optics used to collimate the light source, and absorption and scattering of the sample (ii) shadow artifact originating from a misalignment of the sample center of rotation. (iii) jittered sinogram of a marker bead rotated by a low-cost stepper motor.

### OPT of large organ samples

The non-destructive 3D imaging of whole lung lobes is very useful in the study of COPD models in mice, as it allows the identification of characteristic phenotypes such as bronchoconstriction (narrowing of airways), and the investigation of the extent of the structures affected in different lung areas. The superior, medial, and accessory lobes of the right lung, and the entire left lung of two adult mice were fixed, immunostained, cleared, and imaged using the OptiJ framework (see Supplementary Information for details on mice work). 512 raw projections were acquired over a full rotation of each lobe to obtain high-fidelity reconstructions. Two different proteins expressed in lung epithelial type 2 cells were targeted for fluorescent staining to determine which one allowed for better visualization of the structures critical to studying COPD, such as the bronchioles and alveolar sacs. The lobes of the first mouse were immunostained with a primary antibody against the Surfactant protein C, and the lobes from the second mouse with a primary antibody against the thyroid transcription factor type 1 (TTF-1). In both cases, a secondary antibody conjugated with an Alexa Fluor 488 dye was used to visualize the airway tree through eOPT. The labelling strategy targeting the Surfactant protein C revealed only gross features in the lobes’ eOPT reconstructions, as demonstrated in the orthogonal views of the reconstructed stack from a large left lobe in Fig. 3a-d. The primary bronchus and some secondary and tertiary airways are indicated by red arrows in Fig. 3a-b, and the region in which the indiscernible finer features would be located, the parenchyma (lobe edge), is indicated by red arrowheads. The fluorescent signal collected with this labelling strategy is likely a combination of tissue autofluorescence originating mostly from collagen and the specific fluorescent signal from the dye. The alternative labelling strategy targeting the TTF-1 protein produced reconstructions with an improved signal-to-noise ratio and allowed the visualization of both large airways and minute bronchioles through the center and periphery of the lobes. The orthogonal views of the reconstructed stack from a medial lobe show both the primary and secondary bronchi (red arrows in Fig. 3e-f) and the higher order airways and tiny air sacs in the parenchyma (red arrowheads in Fig. 3.e-f). Figure 3h shows a 3D rendering of the entire medial lobe with a cut-out to direct attention to the intricate network of higher order airways that can be visualized inside the volume. We used Fourier Ring Correlation (FRC) (57) to estimate the resolution of the reconstructed stacks by splitting the data set into two stacks of 256 projections, and obtained a value of 50 μm (see Supplementary Information for details). The reconstructed lung lobes described in Fig. 3 can be viewed and explored interactively using the open-source data visualization platform FPBioimage (53). Volumetric models are available for immersive and interactive viewing directly in standard web browsers at our online repository, along with pre-recorded videos highlighting salient features in the reconstructions: https://lag-opt.github.io.

**Fig. 3.**
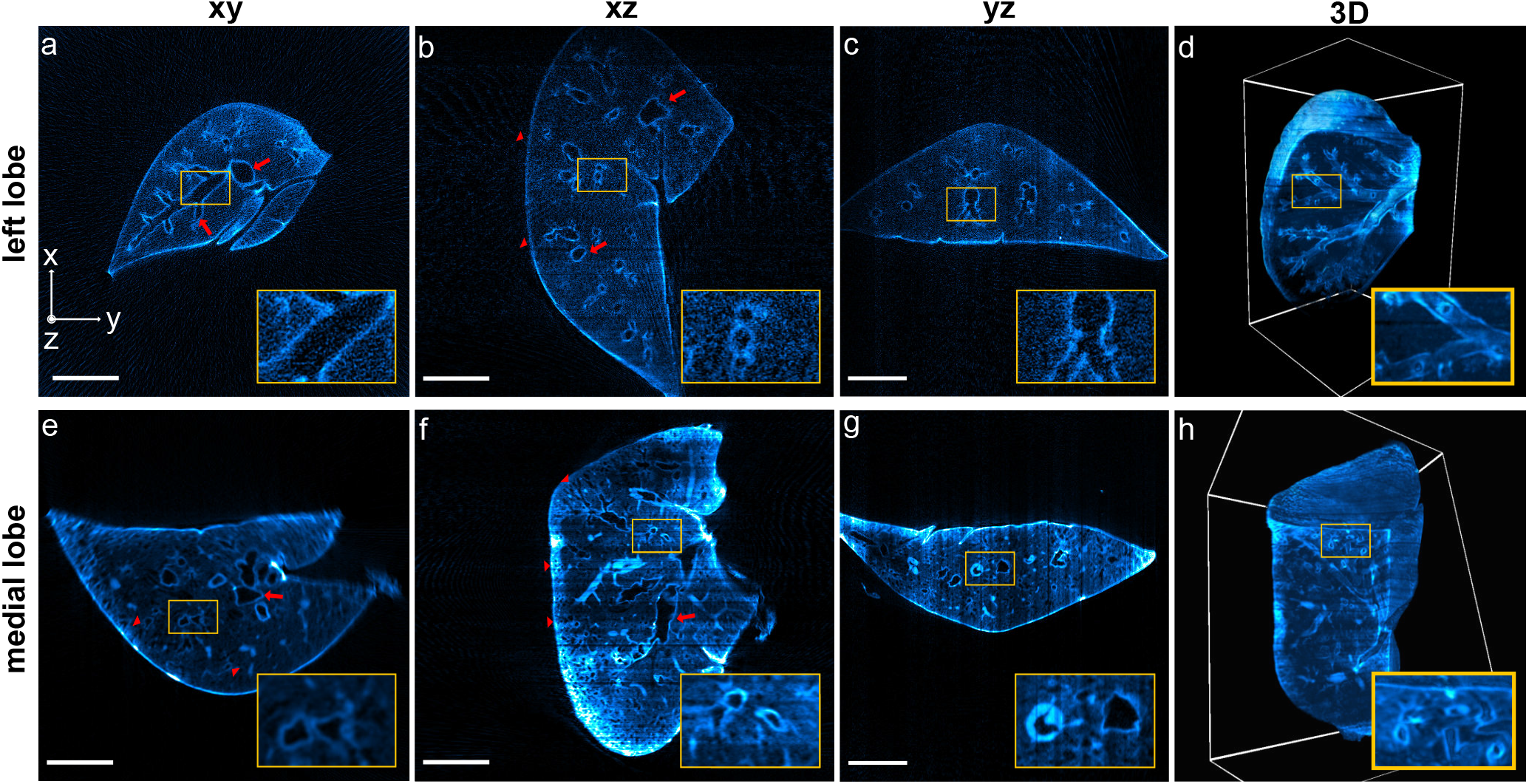
OPT reconstructions of murine lungs. Reconstructions of a left lobe labelled with anti surfactant C – Alexa Fluor 488 (a-d) and a medial lobe labelled with anti TTF1 – Alexa Fluor 488 (e-h) from 512 eOPT projections, displayed in xy, xz, and yz orthogonal views (left three columns), as well as rendered in 3D (right-most column). **a-d)** The red arrows and the insets indicate the primary airways visualized in the orthogonal cross-sections. The 3D rendering in panel **d)** displays a clipping plane through the lung, highlighting secondary and tertiary bronchi in the inset. **e-h)** The red arrows indicate a set of main airways (secondary and tertiary bronchi) in the medial lobe, and red arrowheads indicate high-order airways inside or close to the parenchyma. Small airways close to the primary bronchi are highlighted in the insets on panels **e)** and **f)**. The 3D rendering in panel **h)** with a clipping plane on one of the lobe faces shows a thick meshwork of higher order airways (quaternary bronchi and bronchioles). Interactive 3D renderings are available in our online repository.

## Discussion

OptiJ represents a low-cost open-source hardware and software implementation of OPT for the investigation of large volumetric samples. We demonstrate the imaging of whole organs in 3D with OptiJ at near-cellular resolution. The method reveals the structure of adult murine lungs, from the large primary bronchi to the minute bronchioles at the lung periphery. We compile and provide a novel open-source toolbox of image corrections for OPT measurements and detailed instructions for building a low-cost OPT setup. We present and address the hardware challenges introduced by low-cost OPT solutions. In particular, the sensitivity to sample alignment can be corrected by tracking a marker glass bead and compensating for the tilt using the OptiJ plugins provided. Additionally, we developed a novel Dynamic Offset Correction method to correct for jitter introduced by low-cost stepper motors used for sample rotation. These measures ensure both accuracy and repeatability in the recording of high-fidelity OPT data. Furthermore, we implemented for the first time Fourier Ring Correlation (FRC) as a resolution measure for reconstructed OPT data sets. The non-destructive 3D imaging of COPD mice models lung lobes could provide a whole-organ perspective of alveolar cell clusters in an intact lung, where the involvement of specific cell types in pathophysiological processes could be tracked and quantified, complementary to recent studies of COPD pathophysiology with confocal microscopy (58). Although immunostaining with the anti-surfactant protein C was only partially successful, we were able to make use of autofluorescence from elastin and collagen in epithelial cells and extracellular matrix from the large airway wall to boost signals and obtain high-contrast images of the large airway tree. More generally, the 3D imaging data of intact mouse organs enabled with OptiJ could be useful in tracking specific cell types, visualizing the heterogeneous distribution of disease, or assessing the effects of therapeutics in animal models of COPD. Newer tissue-clearing methods such as 3DISCO (59) and CLARITY (60) can also be implemented to improve on our current approach based on BaBB, which is known to introduce loss of fluorescent signal from certain dyes (61) and may cause linear shrinking of tissue (62). In summary, we provide a unique and complete set of calibration and reconstruction routines in a single ImageJ/Fiji plugin library along with a low-cost, easy to build and easy to use hardware set up. A previous implementation of the Radon transform exists in ImageJ/Fiji, but does not include calibration nor accelerated reconstruction algorithms (63). OptiJ implements both CPU and GPU acceleration for reconstructions, which yields reconstructions in tens of minutes rather than multiple hours. Furthermore, we demonstrate larger fields of view (13×8×8 mm^3^) than most other OPT implementations (17, 22, 24, 28, 38), which typically range from 1×1×1 mm^3^ to 5×5×5 mm^3^. The larger field of view of OptiJ will be useful for examining anatomical structures and fluorescent signals from large model organisms (e.g. mouse, zebrafish, drosophila), organ samples from small animals or even organoids grown from pluripotent stem cells. Future work on OptiJ would include automation of the tilt and offset calibration routines with a direct feedback loop to the hardware after correction with the OptiJ plugins or implementation of deconvolution in OPT data using the model proposed by van der Horst (49). The research presented here was initially conducted in a collaborative effort by a cohort of 14 graduate students and formed part of their PhD training programme in the EPSRC Centre for Doctoral Training in Sensor Technologies and Applications. Students were given a minimal project brief and budget from which they developed a detailed technical proposal and work program. Individuals worked on subsections of the project (e.g. hardware prototyping, software development, biological sample preparation, and data gathering and analysis) with regular supervisory meetings to monitor progress and to identify bottlenecks. The project lasted over a period of 12 weeks and led to the development of a fully functioning prototype of the OPT device presented here. The overall goal was to develop high-end technology that is easily democratised through use of open technologies and open source software and that incentivises further deployment and development by the wider research community.

## Materials and Methods

### Animal ethics

Lung samples were obtained from two naïve C57/Black6 female mice which were humanely euthanised at the end of an independent experiment according to the European ethical guidelines of animal experimentation. The study was approved by the local Ethical committee in Gothenburg (EA137-2014).

### Animal perfusion and tissue preparation

For the immunostaining of the lungs, mice were perfused through the right ventricle with PBS to remove blood from the tissue. Lungs were subsequently inflated with 4% PFA and fixed overnight at room temperature in fixative. Over the next 3 days, the lungs were rinsed in PBS and permeabilised through two cycles of dehydration-rehydration in a gradient of methanol, and in a solution of PBS and detergent (1% Triton X-100) to ensure antigens from the deepest part of the tissue were rendered accessible. All immunostains were then performed in 1% Triton X-100 in PBS (PBST) containing 10% of donkey serum. Two different immunostains were tested in separate lung samples with primary: i) anti-surfactant C protein antibody to target membrane antigen secreted from airway type 2 epithelial cells in alveoli or ii) anti-thyroid transcription factor-1 (TTF-1) antibody (Dako Agilent Products, mouse monoclonal, clone 8G7G3/1, Cat# M3575) to target nuclear antigen also present in airway type 2 epithelial cells. The lungs were incubated in primary antibody solution for 1h at room temperature and for 48h at 4°C followed by extensive washes with PBST and 1% foetal calf serum. Fluorescent labelling of the primary antibody was achieved with anti-IgG Alexa Fluor-488 secondary antibody in 1:500 dilution for 48h at 4°C followed by extensive washes for 3 hr to overnight. A detailed immunostaining protocol is available in the Supplementary Information.

### Sample preparation

Fixed and immunostained samples were embedded in a 2% low-melting-point agarose (Thermofisher Part#R0801) solution as a holding medium for clearing and acquisition. 10 mL syringes were cut using a razor blade at the 1 mL and 6 mL mark. The syringe plunger was inserted from the 6 mL end just so the rubber tip was completely inside the cropped syringe tube. A pipette was used to fill approximately three quarters of the available volume in the tube with molten agarose. The agarose was left to cool for 3-10 minutes, and then samples were carefully transferred into the agarose-filled tube using smooth tweezers and were oriented close to the center of the tube. A spherical glass bead (Sigma-Aldrich Part#Z250465-1PAK) between 0.5 to 1 mm in diameter was immediately inserted close to the sample, but not in the same horizontal plane, as a tracking fiducial for alignment and calibration during postprocessing. The exposed end of the tube was sealed with parafilm to avoid dehydration of the agarose during storage. Samples were placed in a fridge at 4°C for one hour to allow the solution to fully cross-link into solid agarose cylinders. The embedded lung lobes were pushed out of the syringes, dehydrated using 50% methanol for 24 hours and then 100% methanol for 48 hours, and then cleared using a 1:2 mixture of Benzyl alcohol and Benzyl benzoate (BaBB) for 72 hours, changing the BaBB solution every 24 hours. Prior to OPT acquisition, the agarose-embedded tissue cylinders were glued onto bright-zinc plated (BZP) penny washers (M5×25, Fixmart Part#402203217) using quick-dry epoxy (Loctite Epoxy Quick Set 0.85-Fluid Ounce Syringe, Henkel Corporation, Part#1395391). After the glue was cured, the penny washer was coupled to a magnetic kinematic mount (Thorlabs Part#SB1), ready to be inserted into the system for imaging. A detailed description of the preparation and mounting of the murine lung lobes can be found in the Supplementary Information.

### Experimental setup

A 3D-printed flexure stage for opensource microscopy (54), was chosen for x,y,z translation and rotation of the sample because of its low cost (cost of printing material only) and modular design. An Andor CLARA camera with 6.45×6.45 μm^2^ pixels was used for acquisition of the volume projections, although lower cost cameras can also be used. A 0.5x telecentric objective (Edmund Optics Part#63-741) with a 65 mm working distance and 0.028 NA was chosen to acquire the maximum field of view possible with the chosen detector. Two white light LEDs (Thorlabs Part #MWWHD3) were chosen to provide even illumination with minimal flicker. These were fitted in small cage systems with an optical diffuser (Thorlabs Part#DG10-600), an adjustable iris (Thorlabs Part #SM1D12D), and a condenser lens (Thorlabs Part#LA1401-A). A GFP excitation and emission filter pair was used for eOPT (Excitation: 482/25 Part#FF01-482/25-25, Emission: 515/LP Part#FF01-515/LP-25, Semrock). A Hellma glass cuvette (Z805750-1EA, Scientific Laboratory Supplies) was used as the immersion chamber for the sample during imaging. The filled chamber was raised using a Swiss Boy lab jack (Sigma Aldrich Part#2635316-1EA) to completely cover the agarose gel containing the sample during the acquisitions with the BaBB. The acquisition software was written in Java and packaged as an independent executable file. eOPT and tOPT projections were acquired with exposure times of 300 ms and 1 ms, respectively. Pictures of the set-up, a list of parts, instructions for assembly, information about the acquisition software, and the characterization of the *x*, *y*, and *z* motion of the stage can be found in the Supplementary Information.

### Software for image reconstruction

The reconstruction and calibration routines in OptiJ were written in Java and integrated as a plugin library in ImageJ (51), a standard opensource platform for image analysis. OptiJ is available for download online, along with an instruction manual, source code, and examples of use at: https://lag-opt.github.io. The interactive web application FPBioimage was used to visualize the three-dimensional reconstructions of the OPT data used in Fig.3. The reconstructed data sets can be used to visualized and explored online using FPBioimage as well, following the instructions in our online repository.

### Data availability statement

All the raw and processed data, instruction manuals, and code used for this study can be found in our online repository at https://lag-opt.github.io.

## ACKNOWLEDGEMENTS

A functional OPT system prototype was prepared and delivered by the 2015-2016 Sensor Centre for Doctoral Training (CDT) cohort from the University of Cambridge. We thank AstraZeneca PLC for providing dehydrated and stained murine lung samples for imaging and James McGinty and Thomas Watson for fruitful conversations on OPT. We also thank Ricardo Henriques for his neatly formatted BioRxiv preprint LateX template. This work is supported by grants from the UK Engineering and Physical Sciences Research Council, the EPSRC (grants EP/L015889/1 and EP/H018301/1), the Gates Cambridge Scholarship (PVR), the federal government of Nigeria through the Presidential Special Scholarship for Innovation and Development managed by NUC and funded by PTDF (PO), the Wellcome Trust (grants 203249/Z/16/Z and 089703/Z/09/Z), the UK Medical Research Council (MRC) (grants MR/K015850/1 and MR/K02292X/1), MedImmune, the RCUK under the Technology Touching Life Initiative, and Infinitus China Ltd. RFL also acknowledges the support of the UK Biotechnology and Biological Sciences Research Council (BBSRC) TRDF grants (BB/P027431/1 and BB/R021805/1). FS also acknowledges the support from European Molecular Biology Organisation (#7411) and Marie Skłodowska-Curie Actions (#836355).

## AUTHOR CONTRIBUTIONS

P.V.R. did imaging experiments with the mouse lungs, wrote the manuscript, and characterized the OPT system. J.Z. wrote and compiled the suite of calibration and reconstruction routines for the OptiJ software. F-X.B, R.F.L, O.V., F.S., E.J.R., and C.F.K reviewed the manuscript and provided useful feedback. P.V.R. and F-cleared and mounted the murine lungs. O.V. conducted experiments to test early versions of the OptiJ hardware and software components. X-H.Z. perfused and immunostained the murine lungs. F.R. designed and machined the translation and rotation stage for the OptiJ hardware, and designed and built the custom circuit boards used to power and control the LEDs and the stage motors. B.F.S. coordinated the software development, proposed the software tilt and background correction methods, developed the camera interface, and designed the graphical user interface (GUI). P.O. designed, implemented, tested, and packaged the graphical user interface (GUI), and image acquisition software. G.G. made the CAD drawings and wrote assembly instructions for the OptiJ hardware. S.S. tested GPU acceleration in filtered back-projection with preliminary MATLAB scripts. C.V. was the project leader for the Sensor CDT 2015 cohort. O.S. designed the OPT sample holder for 1 mL syringes. F.A. devised the syringe mounting strategy for OPT samples. A.S. prepared early test samples of mice gonads. D.S. characterised the opto-mechanic properties of the OPT, including optical resolution, camera sensitivity and stage positioning errors. F.S. and R.F.L. provided useful advice and helped write software for the OptiJ calibration routines. R.F.L. wrote calibration software in MATLAB to quantify the misalignment of the samples. O.H., R.F.L. and F.S. provided guidance and mentoring for the Sensor CDT 2015-2016 cohort. C.F.K, R.F.L. and F.S. devised the project and organized the Sensor CDT 2015-2016 cohort.

