## Supplementary Information for "OptiJ: Open-source optical projection tomography of large organ samples"

**Vallejo Ramirez et al.**

**Supplementary Figure S1: Left lobe and accessory lobe OPT reconstructions. 2**

**Supplementary Figure S2: OPT resolution estimate using Fourier Ring Correlation (FRC). 3**

**Supplementary Figure S3: Histogram matching. 4**

**Supplementary Figure S4: Sample embedding and mounting for OPT imaging 5**

**Supplementary Figure S5: Diagram and picture of the OptiJ hardware. 6**

**Supplementary Figure S6: CAD Diagrams for OptiJ hardware assembly (steps 1-3). 7**

**Supplementary Figure S7: CAD Diagrams for OptiJ hardware assembly (steps 4-10). 9**

**Supplementary Figure S8: OptiJ acquisition software. 10**

**Supplementary Figure S9: Translation and rotation stage characterization. 11**

**Supplementary Video Legends 13**

**Supplementary Note: Sample clearing and mounting 14**

**Supplementary Note: Mouse lung Immunostaining protocol for OPT 16**

**Supplementary Note: Hardware assembly. 19**

- **Hardware Design 19**
- **Other design considerations 21**
- **Assembly instructions 22**
- **Hardware calibration and characterization 25**

**Supplementary Note: Useful design tips for low-cost OPT 27**

**Supplementary Table S1: Parts List 28**

**References 29**

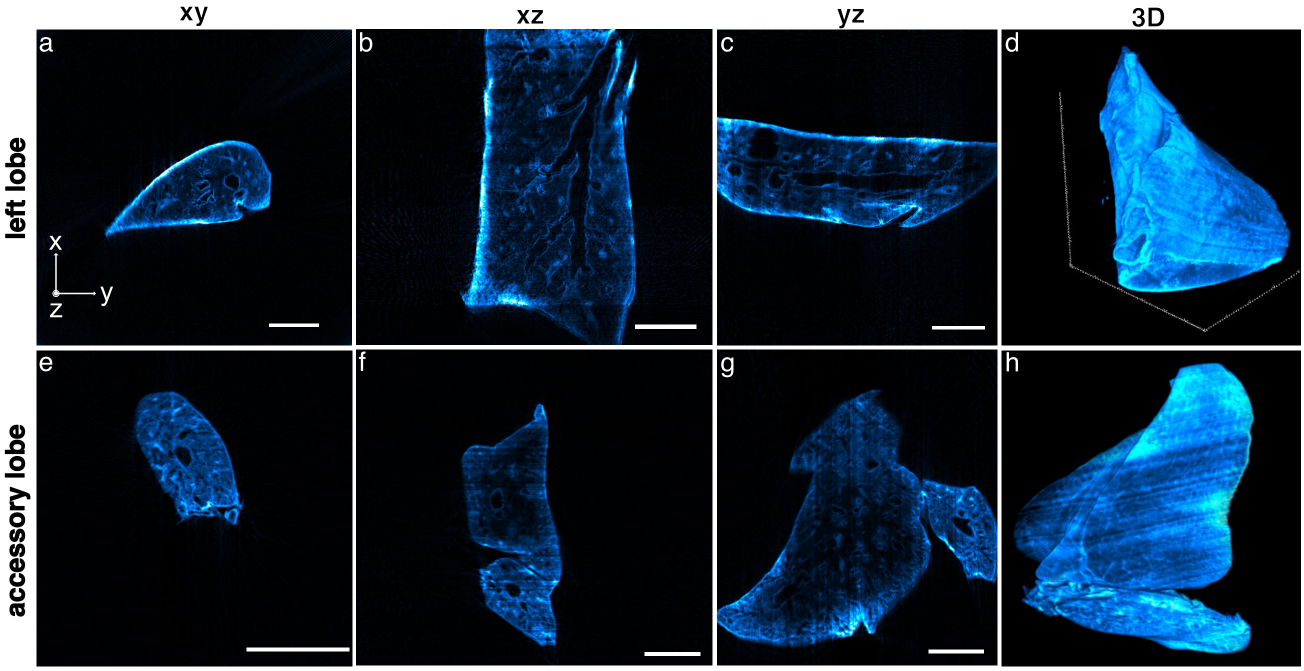


**Supplementary Figure S1: Left lobe and accessory lobe OPT reconstructions.** Volume renderings of two different reconstructed lung lobes are displayed in arbitrary xy, xz, and yz orthogonal views (left three columns), as well as rendered in 3D (right-most column). a-d) Immunostained left lobe reconstructed from 512 raw projections. The 3D rendering in panel h shows the large bottom entrance of the primary bronchus. e-h) Immunostained accessory lobe reconstructed from 512 projections. Scale bar: 2 mm.

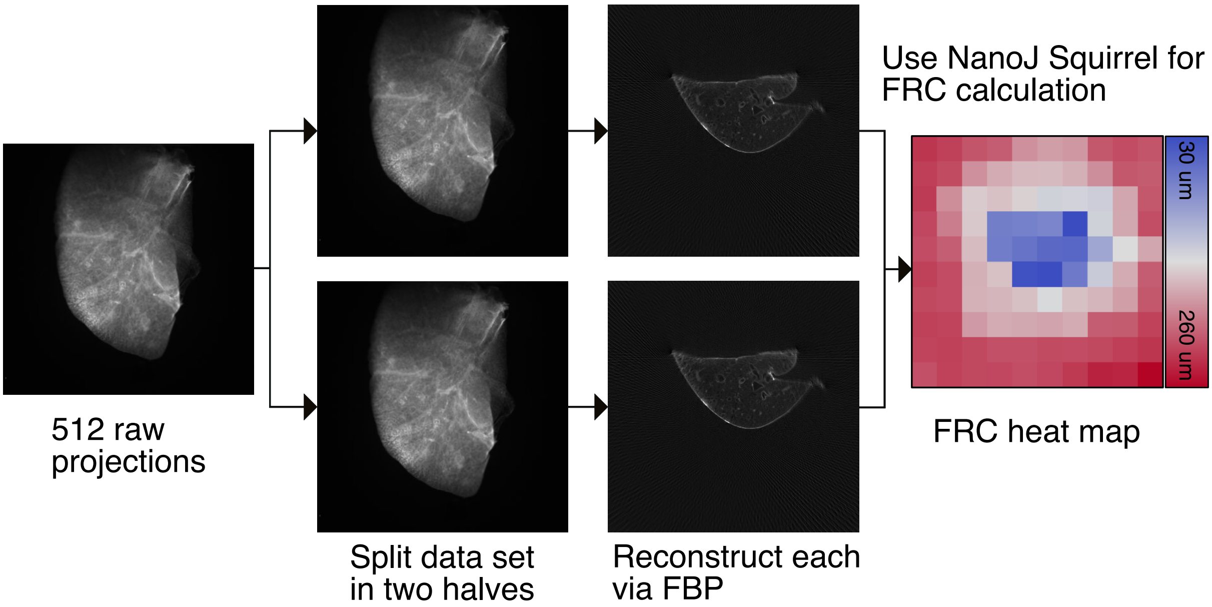


**Supplementary Figure S2: OPT resolution estimate using Fourier Ring Correlation (FRC).** The data set with 512 projections was split into even and odd slice stacks and each half was reconstructed separately using the OptiJ 2D reconstruction plugin. 20 slices were chosen in the middle of the sample and tested with the NanoJ-Squirrel FRC plugin[68]. The reported value of 50 µm for the post-reconstruction resolution of the system is obtained by averaging over the center portion (blue squares) of the FRC heatmaps output by the plugin, representing the pixel area occupied by the reconstructed lung cross-section. The resolution in OPT reconstructions depends on the number of projections, so by effectively down-sampling the data set from 512 to 256 projections, we are substantially underestimating the resolution of the original data set. In the presence of noise and sample jitter, the post-reconstruction resolution of the system is likely to be between 25 and 50 µm.

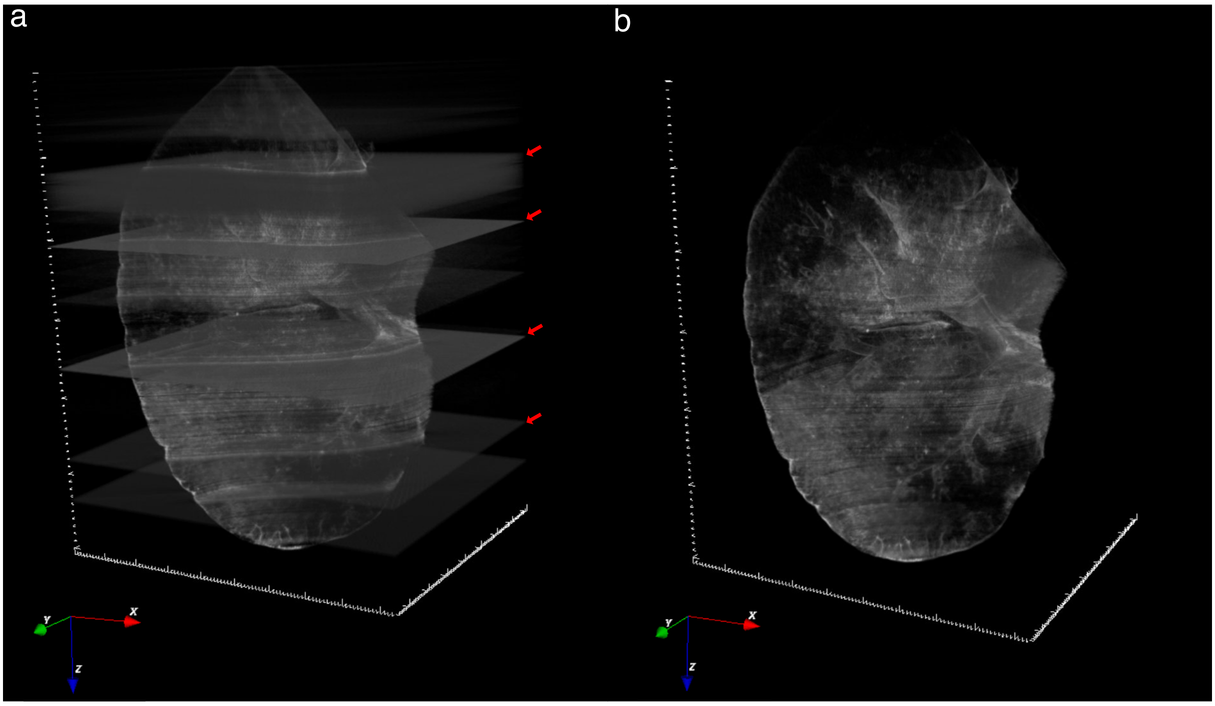


**Supplementary Figure S3: Histogram matching.** eOPT reconstructions may suffer from dark or bright streaks resulting from objects or edges with really strong signals in the sample. These streak artifacts result in very bright or dim cross-sectional slices in the reconstructions, as shown by the red arrows in panel a. We implemented a histogram matching algorithm to account for these intensity variations and we obtain a visualization without bright cross-sectional slices as shown in panel b.

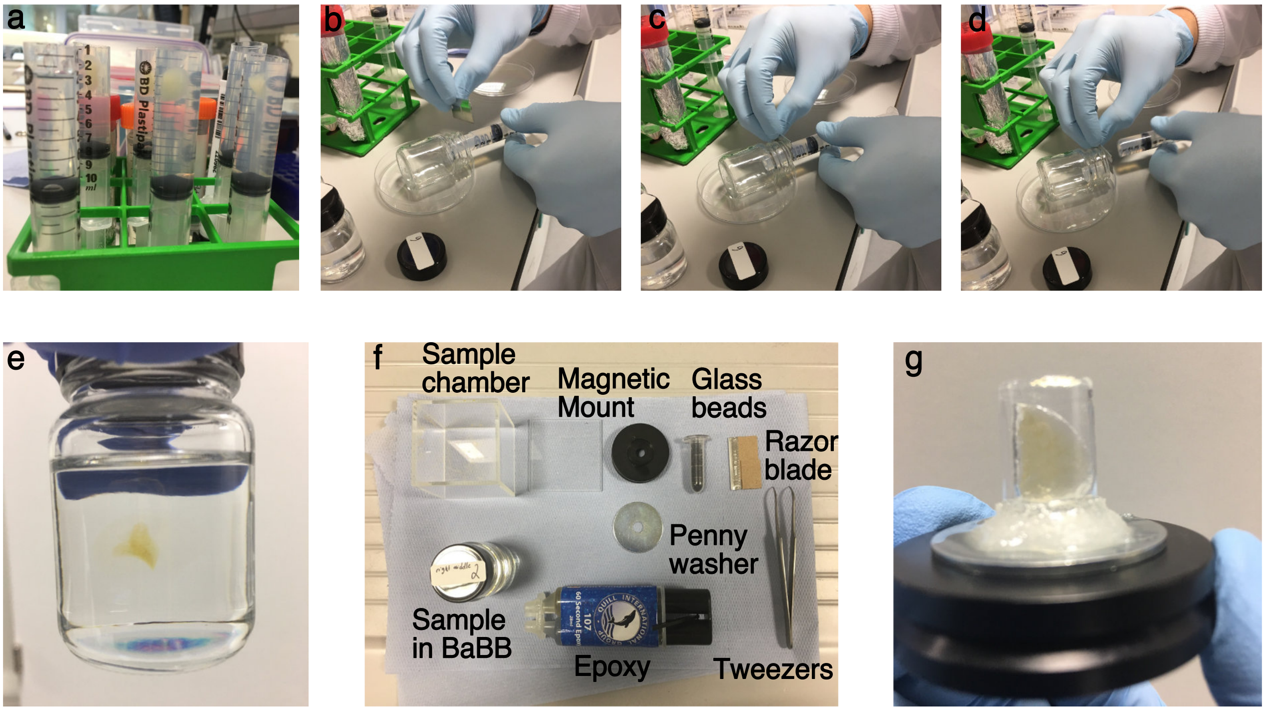


**Supplementary Figure S4: Sample embedding and mounting for OPT imaging** a-c) Embedding of the lung lobes in agarose e-g) Mounting of the cleared lobes on a magnetic stage ready for imaging.

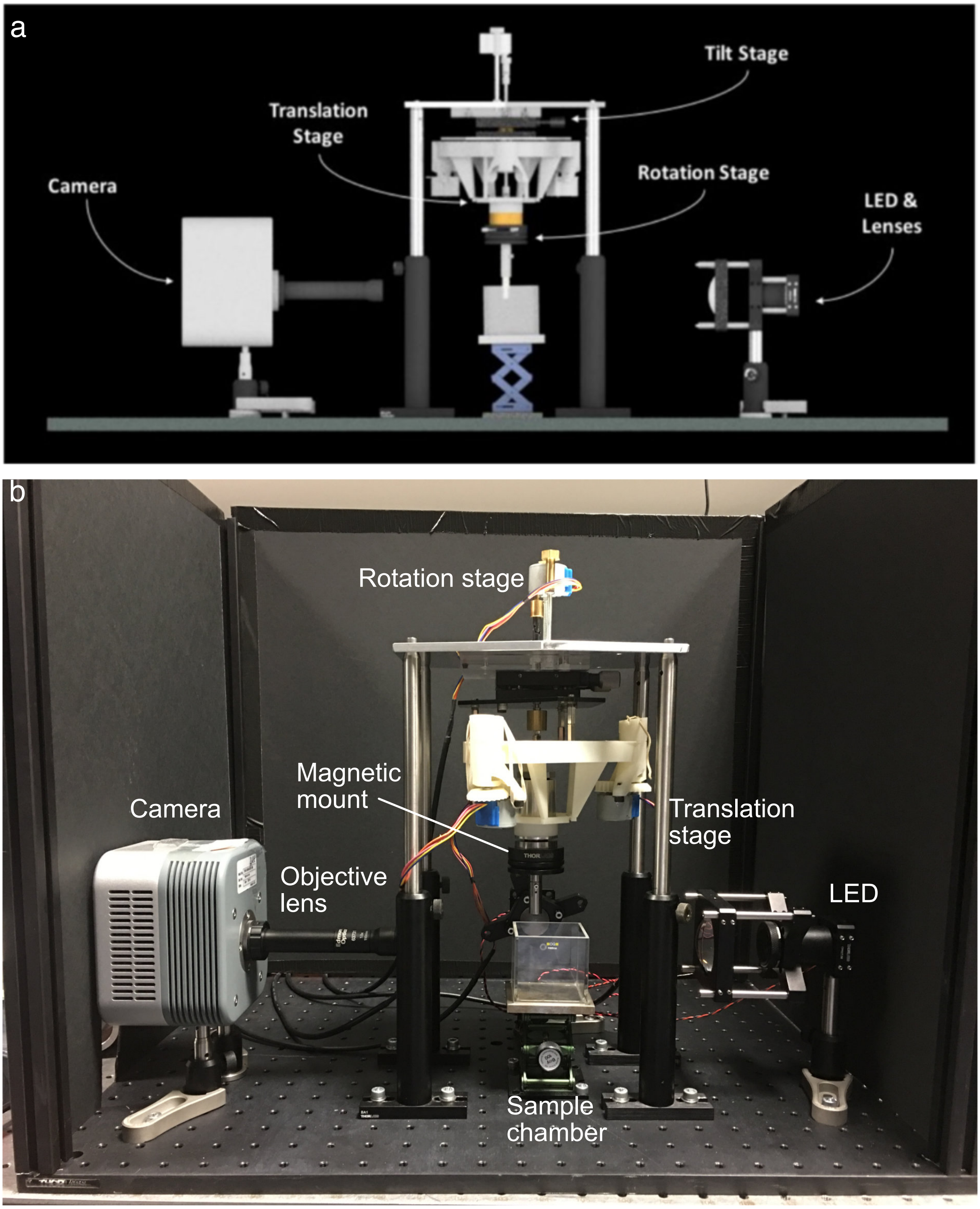


**Supplementary Figure S5: Diagram and picture of the OptiJ hardware.** a) Front-view CAD illustration of the OptiJ hardware. The rotation and translation stages make use of a 3D printed flexure stage that allows for sub-micron precision movement with inexpensive stepper motors. Two LEDs are used for illumination, and off-the-shelf components from Thorlabs are used to support the optics and the stages. b) Picture of the OptiJ setup.

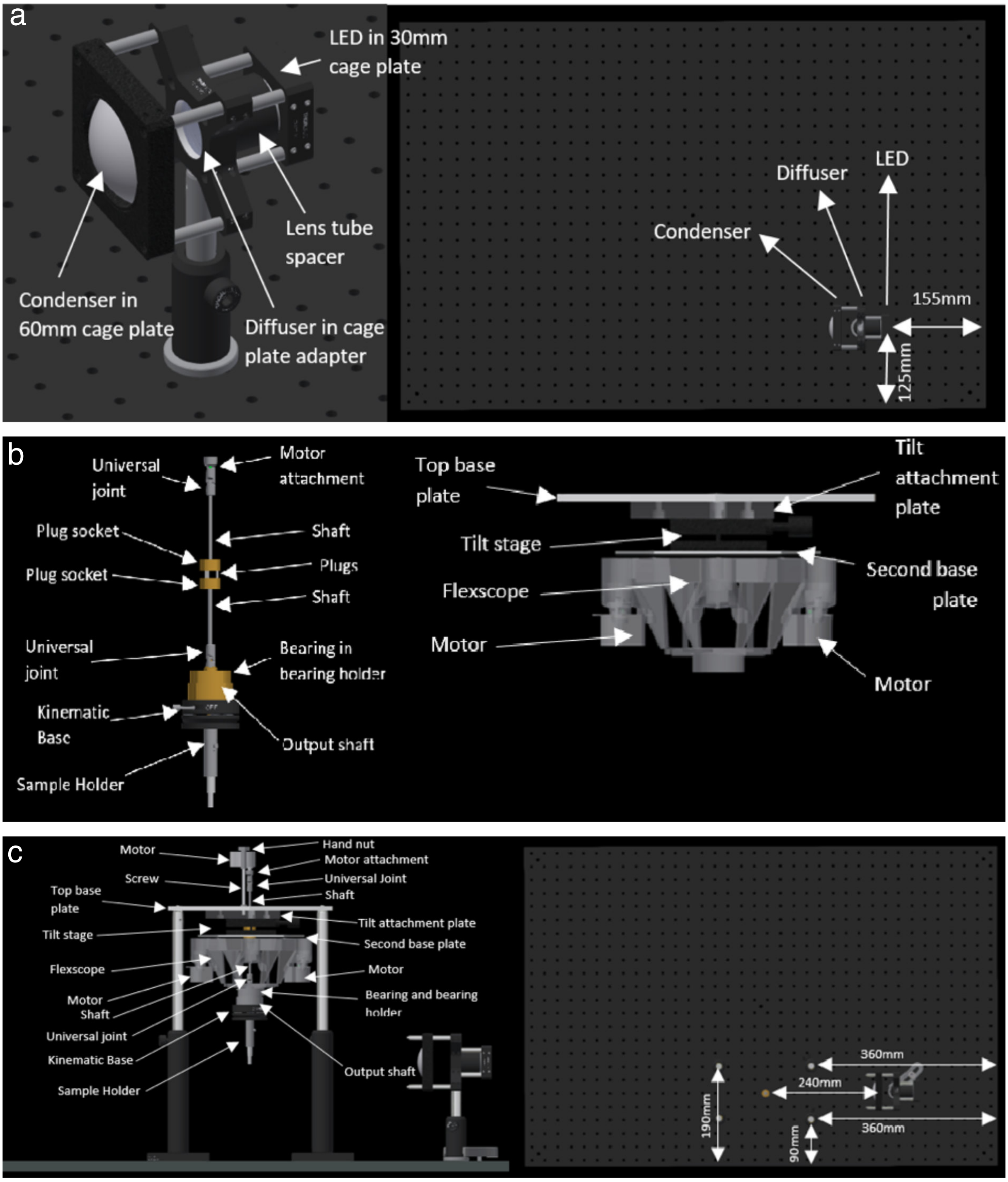


**Supplementary Figure S6: CAD Diagrams for OptiJ hardware assembly (steps 1-3).** a) CAD design of the LED mount with condenser lens and diffuser for transmission OPT using off-the-shelf opto-mechanical components from Thorlabs. The right panel shows the location of the LED on the breadboard with specified dimensions. b) Illustration of the spindle assembly and Deltabot mechanical components. c) Side view of translation and rotation stage assembly using the Deltabot and spindle mounted onto a baseplate. The right panel shows a top-view of the illumination and translation stage assembly on the breadboard with recommended distances.

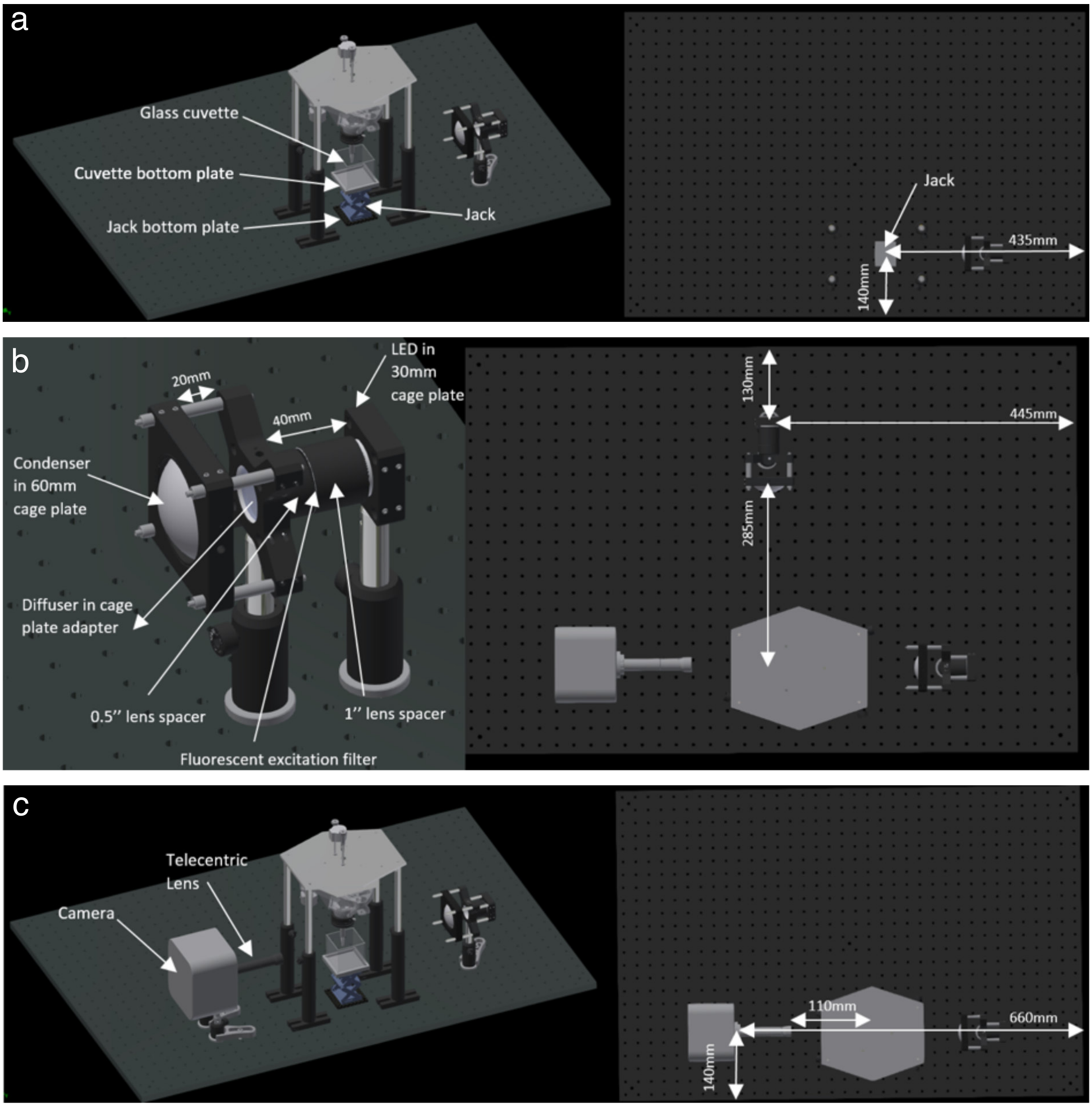


**Supplementary Figure S7: CAD Diagrams for OptiJ hardware assembly (steps 4-10).** a) Perspective CAD snapshot illustrating the position of the Lab Jack under the translation stage. b) The mounting of the LED for fluorescent excitation is similar to that for transmitted illumination, albeit with the addition of a fluorescence filter between the light source and the diffuser. c) Perspective CAD snapshot illustrating the position of the camera relative to the translation stage and the illumination LED

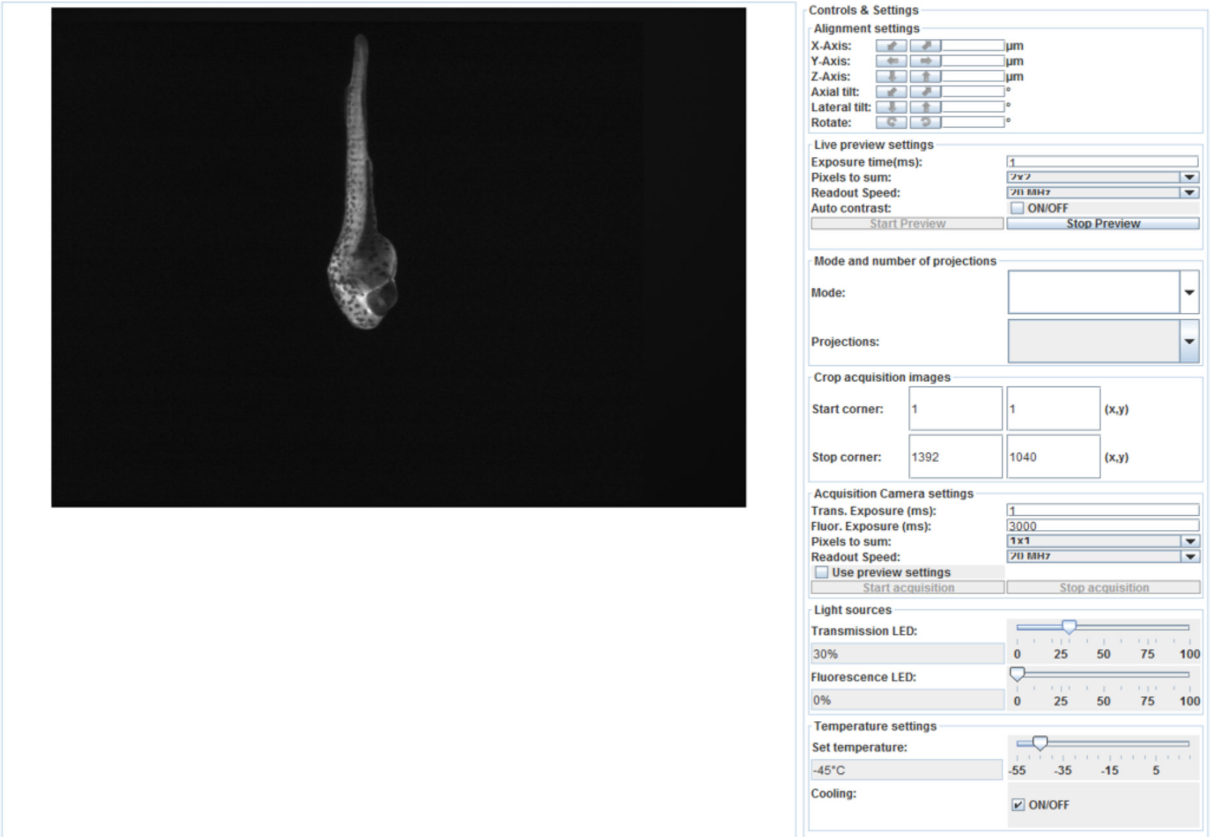


**Supplementary Figure S8: OptiJ acquisition software.** The OptiJ acquisition software is an independent executable written in Java to control the LEDs, motors, and the camera all together via a RaspberryPI®. The GUI includes controls for moving the stage in x,y, and z, and the rotation stage. The GUI includes an option for a live-preview to align the sample in the center of the field of view and to adjust the illumination and imaging parameters. The acquisition parameters enable the user to specify the exposure time, the binning of the camera pixels, and the brightness levels of the tOPT and eOPT LEDs. A screenshot of the software is shown depicting a zebrafish embryo projection in eOPT.

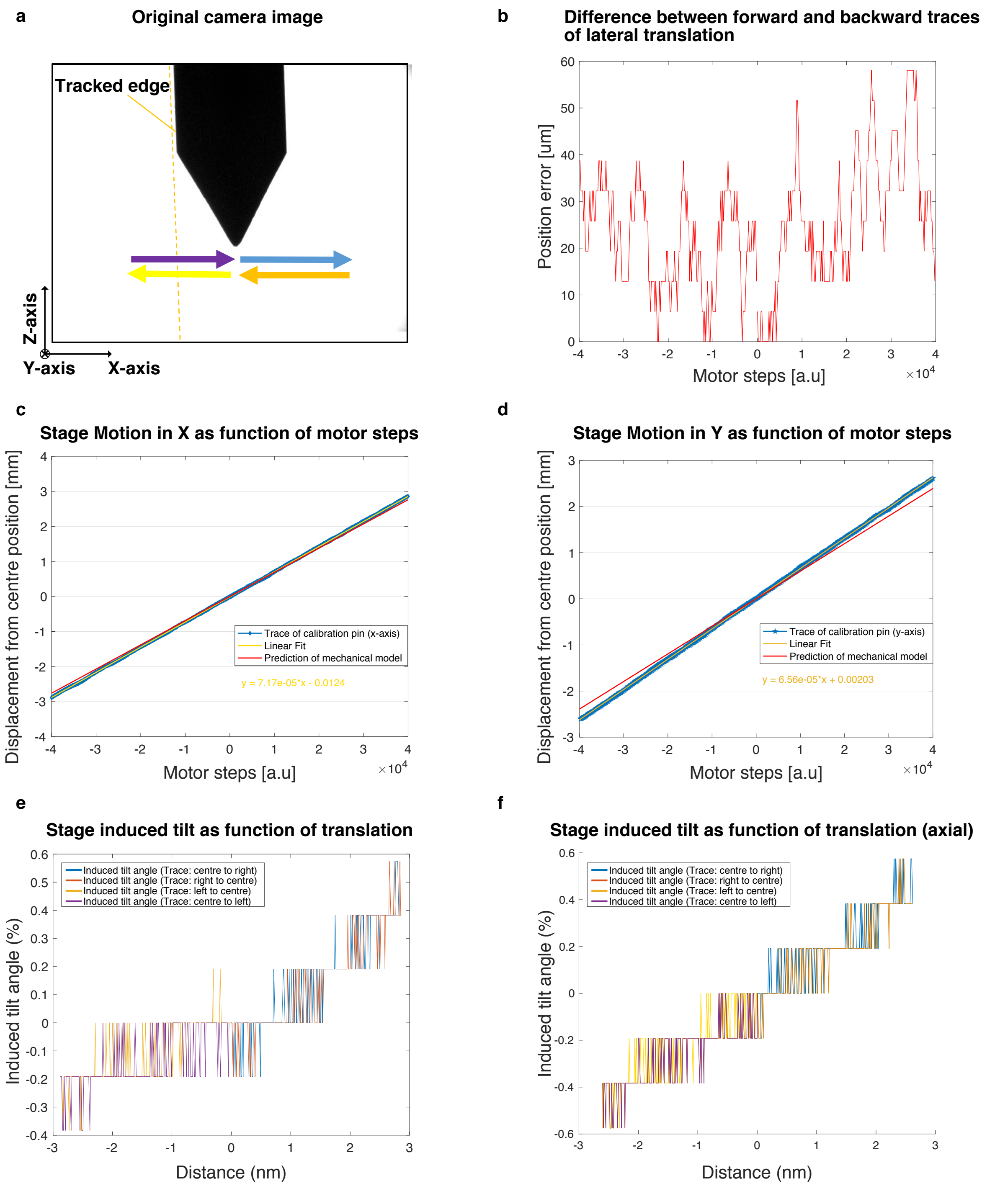


**Supplementary Figure S9: Translation and rotation stage characterization.** (a) Illustration of a calibration pin mounted in place of the sample in the Deltabot. To measure the translation and tilt induced as a function of the motor steps, the pin was moved 3 mm to the right (blue arrow), then to the centre (orange arrow), 3 mm to the left (yellow arrow) and to the starting point again (purple arrow). The side edge on the left of the pin was used to trace translation and induced tilt. (b) The difference between the recorded position of the calibration pin during outwards movement in comparison to its return trace is plotted. Over a travel range of 3 mm in the X direction, the hysteresis of the stage remains less than 58 µm and shows decreased absolute positioning error when staying close to the centre starting point. (c-d) These figures show the motion of the modified Deltabot in the X direction and Y direction, respectively. The data series plotted correspond to the outward and return paths of the calibration pin during the test. From this we confirm the motion is linear, has no significant hysteresis, and corresponds well to the simple kinematic model from the Flexscope design. (e) The relative tilt of the calibration pin in the X direction (lateral tilt) caused by stage motion is shown here across the full range of the stage. The significant quantisation noise in the measurement shows the angular deviation is small on the scale of the system, as it caused only single pixel deviations over the full travel range of the calibration pin. (f) The relative tilt of the calibration pin in the Y direction, termed axial tilt as it occurs along the optical axis, caused by stage motion is shown here across the full range of the stage. As with the lateral tilt measurements, this plot shows that the tilt in this direction is small, linear, and repeatable.

### Supplementary Video Legends

#### Supplementary Video S1

This video shows a left lobe volume with anti surfactant C protein – Alexa Fluor 488 staining. The lobe volume undergoes three full rotations with an opacity change to show the inner structure of the airways inside it. A clipping plane through the mid-section of the lobe shows the large primary bronchus, and a fly-through this airway is shown. The video corresponds to Fig.3.a-d in the main text.

#### Supplementary Video S2

This video shows a medial lobe volume with anti TTF-1 – Alexa Fluor 488 staining. The lobe volume undergoes three full rotations, a fly-through the primary bronchus, and a clipping plane is inserted through its mid-section to reveal the complex airway tree inside. The video corresponds to Fig.3.e-h in the main text. Dark stripes along the volume result from dim cross-sectional slices output by the reconstruction algorithm.

#### Supplementary Video S3

This video shows an accessory lobe volume with anti TTF-1 – Alexa Fluor 488 staining. The lobe volume undergoes three full rotations, and a clipping plane is inserted to reveal the airway mesh inside it. The video corresponds to Fig.S1.e-h.

All videos were created using the open-source web application for 3D data visualization FPBioimage[1].

**Supplementary Note: Sample clearing and mounting**

1. Crop a 10 mL syringe with a razor blade at the 1 mL mark to remove the tapered tip and obtain a plastic cylinder with an open end. Place the syringes on a falcon-tube holder resting on the plunger, with the open side of the cylinder facing the ceiling.

2. Prepare a 2% solution of low-melting-point agarose in milli-Q water and keep the solution in a liquid state by heating it to 60 degrees Celsius.

3. Gently pour the liquid agarose solution into the cropped 10 mL syringes with the plunger at the bottom and let the solution cool for 10 minutes so it begins to form a gel.

4. Pick up the dehydrated and immunostained organ sections with a set of smooth-end tweezers and place them in the middle of the cooling agarose gel, such that the sample remains roughly suspended in the center of the cylinder, with space of 2-5 mm from the open end of the syringe. Let the solution cool and fully cross-link, gently nudging the samples back into place in case they drift towards the sides or the bottom of the syringe. This procedure is shown in Figure S4.a.

a. (Optional) Place a 100 μm glass bead between the sample and the open-end of the syringe, leaving a space between the bottom of the sample and the bead.

5. Fill a set of 200 mL jars with BaBB and a petri dish with milli-Q water.

6. Once the solutions have fully cross-linked and the gel is solid, gently push the end of the agarose block out of the syringe using the plunger, and cut the agarose block using the razor blades, letting the block drop into the petri dish. Gently grab the cylinder with smooth-end tweezers and place it in the BaBB-filled jar. This procedure is shown in Figure S4.b-e.

7. Cover the sample with aluminium foil and place in the fridge for 72 hours, changing the BaBB solution every 24 hours.

8. Remove the agarose block from the BaBB-filled jar using tweezers and place it on a Petri dish. Insert a bead into the agarose block using a set of tweezers if this step wasn't

previously completed in 4.a. Dab a few droplets of epoxy around the center hole of a bright-zinc-plated (BZP) penny washer and quickly place the agarose block in the center of the penny washer, making sure the epoxy holds the edges of the sample in place, as shown in Fig.S4.f Cover with Aluminum foil and let rest for ~30 minutes.

9. Test the bond of the epoxy with the agarose by turning the BZP upside down and seeing if the sample remains in place. Place the BZP penny washer on a magnetic mount.

10. Fill sample chamber with BaBB and couple the magnetic mount to the stage, ready for imaging.

**Supplementary Note: Mouse lung Immunostaining protocol for OPT**

Day 1 (~45min/sample)

1. Perfuse the anaesthetised mouse through the right ventricle with fresh PBS, then with paraformaldehyde (PFA) 4% at room temperature (RT). The lungs are dissected, then inflated with in fresh 4% paraformaldehyde (PFA) at 4°C and immerged in PFA over night with gentle agitation.

Day 2 (2h+ O/N)

2. Flush PBS in the catheter and wash specimens 3x 10 min with PBS at 4°C with gentle agitation,
3. Dehydrate the tissue stepwise in methanol: 33, 66 and 100% for 30 min each.

Note: Tissue can be stored at for a few days at -20°C at this point

4. Rehydrate the tissue back in a series of Methanol 66%, and 33% in PBS for 30min at each step.
5. Incubate in freshly prepared Methanol:DMSO:$H_2O_2$ 30% (2:1:3) at RT with gentle agitation overnight or until the tissue is completely white.

Day 3 (9h+ 48h)

6. Wash specimens for 30 min in 66% Methanol, and 2x30 min in 100% Methanol.
7. Bring the samples to -80°C 4 times for at least 1hr each time and back to RT *(to ensure that antigens in the deeper parts of the tissue are rendered accessible.)*
8. Rehydrate the tissue back to PBST (1% triton X-100 in PBS) in a series of Methanol 66%, 33% and PBST, for 20min at each step.
9. Block nonspecific antibody binding by incubating the lungs in blocking solution (10% of donkey serum =DKBS in PBST 1%) for 48h, changing the bath solution 2 or 3 times during this period.

Day 5(1h+ 48h)

Primary antibodies:

a) Rat anti-protein surfactant C antibody

b) Mouse anti-TTF1 antibody:

Dissolve batch of 0.5mg in 1mL of DMSO than add 9ml of DKBS = 50ug/ml, finally, take 1ml from last dilution and add 9 mL of DKBS= 5 ug/ml.

10. Incubate the tissue for 1 hr in primary antibody (Rat anti-protein surfactant C mAb or Mouse anti-TTF1 mAb) diluted in blocking solution (DKBS) at RT and then for 48hrs at 4°C, with gentle agitation.

Day 7 (3h+O/N)

11. Rinse thoroughly the samples in PBST+1% HI-FCS as follows, with gentle agitation:
    - 1x1ml 1hr at RT
    - 1x4ml 1hr at RT (in glass vials)
    - 3x4ml 1hr at RT (in glass vials)
    - O/N at RT (in glass vials).

    *Thorough rinsing is crucial to ensure that unbound antibodies do not remain trapped within the tissue and give an unacceptably high background staining.*

Day 8 (1h+48h)

12. Incubate the tissue for 48h at 4°C in secondary antibody (Alexa 488 anti-mouse IgG) diluted in blocking solution. Stock solution (0.4 mg/ml in Glycerol:H2O 1:1 in the freezer) to be diluted to the 1:1000 in blocking solution. Pipet 15 µL of the stock solution and add to 15 mL of blocking solution.

Day 10 (3h+O/N)

13. Once again rinse thoroughly the samples in PBST (without HI-FCS) as follows, with gentle agitation:
    - 1x1ml 1hr at RT
    - 1x4ml 1hr at RT (in glass vials)
    - 3x4ml 1hr at RT (in glass vials)
    - O/N at RT (in glass vials).

    *Thorough rinsing is crucial to ensure that unbound antibodies do not remain trapped within the tissue and give an unacceptably high background staining.*

Day 11 (3h+ 72h)

14. Mount the tissue in 1% low melting agarose (dissolved in milli-Q water)

15. Dehydrate the agarose-embedded tissue in 50% methanol for 24h and then 100% (Methanol) for 48hrs

Day 14 (30min+ 72h)

16. Clear the specimen in BABB solution (1:2 benzyle alcohol, benzile benzoate) for 72hrs changing the solution every day.
17. Scan in OPT. The specimen can be stored in BABB (in the dark) for up to several weeks.

**Supplementary Note: Hardware assembly.**

### Hardware Design

A 3D printed flexure stage for open-source microscopy[2] was chosen to hold the sample and move it in x, y, and z because of its low cost (cost of printing material only) and modular design. The design of the stage was modified into a triangular geometry to accommodate for a rotation arm through the center of the structure. It was fitted with cost-effective stepper motors that move a set of gears connected to actuators to induce motion in three orthogonal directions.

The stepper motor used for rotation was observed to introduce motion jitter during acquisitions. This causes artifacts during reconstructions, as data from multiple horizontal planes from the sample maps onto the same row of pixels. A custom spindle with high-precision angular contact bearings was designed and machined to allow for a smooth transfer of motion between the stepper motors and the sample holder. An ANDOR Clara camera was first implemented as a detector as it was readily available—however a less expensive option with the similar specifications can be used instead. The ATIK 414EX and ATIK420 cameras were tested with OptiJ and can be readily implemented with similar performance. Warm white LEDs from Thorlabs were chosen to provide even illumination with minimal flicker (to avoid differences in light levels from one projection to the next as the sample rotates) and a broad excitation spectrum to be compatible with different fluorescent dyes. The rest of the hardware is composed of standard off-the-shelf opto-mechanics from Thorlabs to hold the components on an optical breadboard.

Three custom printed circuit boards (PCBs) were used to power and control the motors for x,y,z motion and rotation, as well as the light-emitting diodes (LEDs) for illumination. These were chosen to fit the design of the Deltabot and the four motors it uses. A full list of parts and detailed instructions on how to assemble them can be found in the following section. The design of this OPT system followed an implementation similar to Sharpe[3] and Henkelman[4]. Figure S5.a shows a CAD drawing of the OptiJ setup, and Fig.S4.b shows a picture of the set up with labelled components. The illumination is set up in two separate arms, one path along the optical axis for tOPT and a path orthogonal to the optical axis for eOPT. The sample is held in place using a magnetic mount coupled to the Deltabot, which is mounted upside-down onto a metallic plate fastened to four tall optical posts. The camera is mounted in line of the optical axis. The sample chamber (filled with immersion fluid during acquisitions) is placed on top of a small lab jack so it can be raised and lowered to cover the sample.

### Other design considerations

Unlike many forms of microscopy, the resolution obtained in an OPT implementation is contingent primarily on high-fidelity motion control in full 3D. In our implementation, the motion control of the stage was partitioned into mechanically distinct elements, one to achieve translation and two separate rotation stages. The translation stage selected was adapted from the open source 'flexscope' project[2], which offered design flexibility and low cost of implementation through a one-piece 3D printed construction. A rotation stage was included to enable the sample rotation required for OPT. For this purpose, a simple spindle was designed and fitted with high precision ABEC9 angular contact bearings, with the aim of minimising sample jitter during rotation. The second mechanical correction component was a commercially available tilt stage to correct for axial tilt; rotational displacement of the sample's rotation axis from alignment with the image plane caused by real-world misfits and misalignments, as well as unintended rotation caused by the translation stage. Acting together, these three stages enable the operator to position the sample in the centre of the image plane, corrected for angular misalignment and complete an imaging cycle. Automation of the translation and sample rotation stages was accomplished using low cost stepper motors (Part #28-byj48), connected to the computer running the imaging software through open-source motor drivers, which offer a simple interface to the imaging software and implementation flexibility due to their modular design.

Another important consideration for OPT is temporal and spatial illumination uniformity. Warm white LEDs from Thorlabs were chosen to provide even illumination with minimal flicker, which improves image quality by minimising differences in light levels from one projection to the next as the sample rotates. Illumination intensity was controlled by a modified motor driver, reconfigured to allow precise control the current supplied to the LED, which exploited the flexibility of the custom-built electronics architecture to allow for seamless control of both sample positioning and illumination from the same interface.

### Assembly instructions

1. Mount the LED (Part MWWHD3) in a 30 mm thick cage plate (Part CP02T/M) and attach a 1 inch lens tube spacer (Part SM1S10) while also placing four 6 mm diameter cage rods within the cage system. Mount the diffuser (Part DG10-600) in a 30-60 mm cage system adapter (Part LCP02-M) and attach to the 30 mm cage plate using the rods. Attach the cage system adapter to an optical post (Part PH50/M and TR50/M) and place on the breadboard. Align the optical post so that the centre of the post is approximately 170 mm from the right-hand short edge of breadboard and 140 mm from the bottom long edge. Mount the condenser (Part LA1401-A) in a 60mm cage plate (LCP01/M) and attach at a distance of 15mm from the cage plate adapter using 4x6 mm diameter cage rods.

2. The first step is to create the motor bearing system. Place the 3 plugs into the top plug socket then attach it to the bottom plug socket. Screw in the two shafts into both the top and bottom plug socket. Glue each shaft into a universal joint and attach the top universal joint to the motor attachment. Attach the tilt stage (Part 55-459) to the tilt attachment plate using two 4 mm screws on either side of the central hole of the tilt stage. Attach the tilt stage to second base plate using the four 4 mm attachments in each corner. Attach the second base plate to FlexScope using the three 4 mm attachments. Attach the tilt attachment plate with the previously described parts already mounted to the top base plate using the three 4 mm attachments. Then place the motor bearing system through the middle holes and attach the motor attachment to a motor through the top base plate with two screws and the hand nuts. Join the bottom universal joint to the output shaft with the bearing holder with bearing within and the bottom part of the Flexscope between them. Join kinematic base to bottom of output shaft and attach sample holder to kinematic base. The completed assembly is shown in Fig.S6.a.

3. Attach the top base plate to 4 large optical posts (Part PH50/M and TR150/M) using the four external 4 mm taps in the corners of the top base plate and mount on the optical breadboard so that the sample holder is central to the central line of the LED and located 240 mm from the closest edge of the condenser lens mount. The two posts closest to the transmission LED will be placed approximately 90 mm and 190 mm from the bottom long edge and 360 mm from the right-hand edge as shown in Fig.S6.c.

4. Attach the bottom of the lab jack (Part 2635316-1EA) to the breadboard under the centre of the sample holder, ensuring that the edge of the jack is parallel to the objective lens. The centre of the jack will be approximately 435 mm from the right edge of the breadboard and 140 mm from the bottom edge of the breadboard. Attach the cuvette bottom plate to the top of the jack and place the glass cuvette (Part Z805750-1EA) on top, as illustrated in Fig.S7.a.

5. Attach the telecentric lens (Part #63-731) to the camera using the appropriate mount or converter. Mount the camera on two optical posts along the central line so that the end of the telecentric lens is sitting 110 mm from the centre of the sample holder. The camera edge attached to the telecentric lens will be approximately 660 mm from the breadboard right-hand edge and the centre of the telecentric attachment will be 140 mm from the bottom long edge of the breadboard.

6. Activate the live video mode with the camera's native software to align the illumination optics.

7. Activate LED illumination using a RaspberryPI, Arduino, or other interface to control the electronics.

8. Adjust the height of the LED so that the center of the illumination spot coincides with the center of the camera field of view.

9. Adjust the distance and height of the condenser relative to the diffuser until the illumination in the camera's active area is roughly uniform.

10. Attach the second LED (Part MWWHD3) to a 30 mm thick cage plate (Part CP02T/M). Attach the cage plate to the 1 inch lens spacer (Part SM1S10). Place the fluorescent excitation filter (Part 84-101) between the 1 inch lens spacer and the 0.5-inch lens spacer (Part SM2L05). Mount the LED cage plate to an optical post (Part PH50/M and TR50/M) and place on breadboard 445 mm from right-hand edge and 130 mm from top long edge, as shown in Fig.S7.b.

11. Mount diffuser (Part DG10-600) in a 30 mm to 60 mm cage system adapter (Part LCP02-M). Place on optical post (Part PH50/M and TR50/M) and attach optical post to breadboard 450 mm from right hand edge and 185 mm from back edge. Mount condenser in 60 mm cage plate (LCP01/M) and attach to cage plate adapter using 4 x 6 mm diameter cage rods 20 mm from the diffuser cage plate.

12. Turn on the fluorescence LED and use the two cage alignment plates to ensure that the LED is focussed on the center of the cuvette (place a piece of paper inside the cuvette to observe the excitation spot).

13. Place a piece of white paper at the level of the alignment pin and move the fluorescence LED backwards and forwards but not sideways to create even illumination on white paper where the sample will be placed.

14. (Optional) Open the OptiJ acquisition software (independent executable written in Java, and set up acquisition parameters.

### Hardware calibration and characterization

As the Deltabot forms the basis for the 3D positioning capabilities of the stage, it is important that the relationship between its input and output is well understood. The motors that actuate the stage are controlled by electronic motor drivers, which were assumed in this particular analysis to operate perfectly without missing steps. As such, the input to the Deltabot was taken as the number of steps sent to the motor controller, and the output as the motion of a calibration pin as indicated in Fig.S9.a. As motion in the Z-axis was considered to be less critical to the overall operation of the system than motion in X and Y, the latter two axes being altered on a more frequent basis during regular operation, analysis of X and Y motion only is presented here for brevity.

Rotation of the stage platform relative to its starting position is referred to as 'tilt' in this analysis. Measuring the tilt is important as it violates the assumptions of the analysis used to reconstruct images and therefore degrades image quality. Figure S.9.e-f show the result of measuring tilt due to stage motion, demonstrating that not only is the tilt relatively small - reaching a maximum of 0.6 degrees over a travel range of 3 mm – but also shows linear repeatable behaviour.

In Fig.S9.c-d the relationships between the number of motor steps and the motion in X and Y direction are shown. Both plots show a strong linear relationship between motor steps and stage motion over a wide translation range that shows negligible dependency on the motion history of the stage. Additionally, the stage was found to exhibit no measurable time dependencies on the timescales of interest (1-1000 s) through tests in which multiple images of each position were captured. Moreover, the difference in absolute positioning between a forward and backward trace is shown in Fig.S9.b. For a whole round trip of the calibration pin the backward scan deviates up to 58 μm from the first trace. The coefficient C, relating motor steps to millimetres of motion as derived from the linear least squares fit of the data in each plot, is of particular interest as it can be predicted from the geometry of the stage. The theoretical relationship is as follows:

$$Cx = Lever\ Ratio\ \times Gear\ Ratio \times \frac{Thread\ Pitch}{Steps\ per\ Revolution}$$

for the X direction and

$$Cy = \frac{\sqrt{3}}{2}Cx$$

for the Y direction. The factor of $\frac{\sqrt{3}}{2}$ which relates the two coefficients, derives from the basic geometry of the Deltabot, in which each actuator moves the platform along a line that is at a 30 degree angle to the lines that join the actuators. Thus, as the X direction in the system is aligned with one of the actuators' directions of motion, the Y direction is at a 30 degree angle to the remaining two actuators' directions of motion. In the case of the stage design used in this system, *Lever Ratio* is the ratio of the stage height to the length of the actuator arm. For the gears used in this system, the *Gear Ratio* is fixed at 2. The closeness of the predicted value for the coefficients *C* to the measured value is shown by the alignment of the “Prediction” dataset with the measured data in the figures, which show that *Cx* was measured at 3.77% more than the predicted value, and that *Cy* was measured at 9.00% more than the predicted value. This discrepancy cannot be entirely accounted for by defects in the dimensions of the printed stages, as the stages were measured to be accurately printed to within 0.5 mm, nor can it be accounted for by a misalignment in the system's axes with the camera axes, which would cause an equal decrease in the measured value from the predicted coefficient value for both coefficients. It is therefore suggested that the discrepancy is due to motion derived from other mechanisms than are not included in the simple linkage analysis presented here, though it should be noted that as the behaviour of the stage is both linear and stable, it is quite sufficient in many applications simply to use the stage as a black box mechanism.

**Supplementary Note: Useful design tips for low-cost OPT**

Adhesive for sample mounting

We tested two different adhesives to couple the agarose cylinders containing the sample to the penny washers that magnetically attach to the translation and rotation stage: quick-dry epoxy and super glue (cyanoacrylate). Both adhesives were gently poured on top of the penny washer and the agarose cylinders were gently pressed on top to immerse one end of the cylinder in the adhesive. After letting the adhesive cure with the sample upside down, as shown in Fig.S1.g, the sample was immersed in BaBB for imaging. The samples bonded with cyanoacrylate became cloudy, as the BaBB either partially dissolved or reacted with the cyanoacrylate, contaminating the index-matched medium. This cloudy effect was not observed with the epoxy, even after immersion for several hours. We therefore used quick-dry epoxy for the remainder of our experiments.

Sample drift from epoxy dissolution

The agarose cylinders in which the sample is mounted may drift towards the bottom of the camera field of view during long acquisitions, as the BABB solution partly dissolves the epoxy used to glue the cylinder to the BZP penny washer. The drift is typically small (a few pixels) and can be corrected by tracking a marker bead or fiducial in the sample.

Motion jitter from cheap stepper motors

Using cheap motors for rotation can lead to significant sample jitter during acquisitions (which can be identified as sharp edges in an otherwise smooth sinogram). This jitter can be addressed first using the *Dynamic Offset Correction* routine in OptiJ, but it can also be corrected using more expensive motors or ball bearings to smoothen out any jitter from the cheap motors.

| Part | Supplier | Part Number | Quantity | Price (GBP) |
|---|---|---|---|---|
| Condenser Lens | Thor Labs | LA1401-A | 2 | 32.63 |
| Diffuser | Thor Labs | DG10-600 | 2 | 10.73 |
| Cage Plate Adaptor | Thor Labs | LCP02-M | 2 | 27.93 |
| Thick Threaded Cage Plate | Thor Labs | CP02T/M | 2 | 15.44 |
| Large Cage Plate | Thor Labs | LCP01/M | 2 | 27.27 |
| Lens Tube Spacer 1 | Thor Labs | SM1L05 | 1 | 9.25 |
| Lens Tube Spacer 2 | Thor Labs | SM1S10 | 2 | 8.82 |
| Cage Rods | Thor Labs | ER2-P4 | 16 | 16.59 |
| Short Posts | Thor Labs | TR30/M | 2 | 3.48 |
| Medium Posts | Thor Labs | TR50/M | 2 | 3.81 |
| Long Posts | Thor Labs | TR300/M | 4 | 7.95 |
| Short Post Holders | Thor Labs | PH50/M | 4 | 5.66 |
| Long Post Holders | Thor Labs | PH150/M | 4 | 9.29 |
| Optical Breadboard | Thor Labs | MB6090/M | 1 | 490.25 |
| Post Holder Clamps | Thor Labs | CF175 /M | 8 | 8.45 |
| LED | Thor Labs | MWWHD3 | 2 | 9.00 |
| Objective Lens | Edmund Optics | 63-731 | 1 | 396.00 |
| Filter Mount | Edmund Optics | 65-800 | 1 | 28.00 |
| Filter Adapter | Edmund Optics | 88-213 | 1 | 23.60 |
| Excitation filter | Semrock | FF01-482/25-25 | 1 | 350.81 |
| Emission filter | Semrock | FF01-515/LP-25 | 1 | 407.72 |
| **Mechanical Stage** | | | | |
| Tilt Correction Stage | Edmund Optics | 55-459 | 1 | 207.20 |
| Stepper Motors | Ebay | 28BYJ-48 | 5 | 1.86 |
| Cuvette | Scientifc Laboratory Supplies | Z805750-1EA | 1 | 83.84 |
| Lab Jack | Sigma-Aldrich | 2635316-1EA | 1 | 69.90 |
| Rotation Mechanism | | | | |
| Stepper Motors | Ebay | 28BYJ-48 | 1 | 1.86 |
| Universal Joints | RS | 689-215 | 2 | 69.24 |
| Bearing | Quality Bearings Online | 7000A5TRSULP3 | 1 | 69.47 |
| Kinematic Stage | Thor Labs | SB1/M | 1 | 70.27 |
| **Electronics & Control** | | | | |
| Raspberry Pi | RS | 832-6274 | 1 | 27.99 |
| Motor Driver Boards | Newbury Electronics | | 1 | 101.58 |
| LED Driver Boards | Newbury Electronics | | 1 | 187.94 |
| Driver Board | Components Farnell | | | 260.28 |

**Supplementary Table S1: Parts List**
